# Identifying brain areas for tactile perception through natural touch with MR-compatible tactile stimulus delivery system

**DOI:** 10.1101/2022.10.24.513441

**Authors:** Seong-Hwan Hwang, Doyoung Park, Somang Paeng, Sue-Hyun Lee, Hyoung F. Kim

## Abstract

Primates actively touch objects with their hands to collect information. In investigations of the tactile information processes, participants should experience tactile stimuli through active touch while brain activities are monitored. Here, we developed a pneumatic tactile stimulus delivery system (pTDS) that delivers various tactile stimuli on a programmed schedule and allows tactile perception through voluntary finger touches during MRI scanning. A pneumatic actuator moved tactile blocks and placed one in a finger hole. The time when an index finger touched a tactile stimulus was detected with a photosensor, allowing analysis of the touch-elicited brain responses. The brain responses were examined while the participants actively touched braille objects presented by the pTDS. BOLD responses during tactile perception were significantly stronger in a finger touch area of the contralateral somatosensory cortex compared with that of visual perception. This pTDS enables MR study of brain mechanisms for tactile processes through natural finger touch.

## Introduction

In daily life, primates including humans process multiple sensory information received from the external environment. In particular, the processing of tactile information that is obtained through direct skin contact provides useful information immediately. For example, whether an object is an apple or an orange and whether it is fresh or not can be distinguish by touch. Through the perception of tactile stimuli, primates can recognize object identity, comprehend the current condition, and make an appropriate response to the objects. Understanding how tactile perception is processed in the brain is therefore fundamental to understand perception and behavior.

While visual perception research occupies the majority in cognitive neuroscience, some studies have focused on tactile perception research. To examine neural responses during tactile perception in the primate brain, researchers have attempted to develop devices for presenting tactile stimuli in the Magnetic Resonance (MR) scanner (Debowska et al., 2013; Hegner et al., 2010; Kitada et al., 2019; Lee et al., 2019; Podrebarac et al., 2014; Zhang et al., 2005). Most devices were designed to present a tactile stimulus to the fingers of humans whose movements were restricted (Hegner et al., 2010; Kitada et al., 2019; Zhang et al., 2005). In this restrained condition, however, it is difficult to test naturalistic tactile perception by active finger movements. Because tactile perceptions mostly occur through finger and hand movements, there may be a gap between tactile processing in the restrained state and in the natural state. To investigate naturalistic tactile perception processed in the human brain, it is necessary to allow active touch by finger movements during fMRI research.

Allowing finger and hand movements in MR scanner is challenging because the movement can produce movement-induced noises including motion noise and brain responses unrelated to tactile perception (Middlebrooks et al., 2017). The brain responses induced by tactile stimuli must be identified, and separated from movement-induced noises. Moreover, to accurately examine the brain response elicited by actively touching a tactile stimulus, it is necessary to detect the timing when the finger touches a tactile stimulus during the fMRI experiment.

Furthermore, there are various properties of tactile stimuli such as texture, shape, softness, and stickiness (Hegner et al., 2010; Kitada et al., 2019; Lee et al., 2019; Podrebarac et al., 2014; Zhang et al., 2005). These various properties can be recognized and the information used for survival. Therefore, a device that can test different tactile properties is essential to investigate the brain processes of different tactile information. So far, however, how to physically deliver different types of tactile stimuli for investigating the active tactile perception in an MR scanner has not been established. In addition, a programmable system for delivering the stimuli is required to coordinate the presentation of various tactile and visual stimuli together in one task. This led us to design a device in which an experimenter outside the scanner plans which tactile and visual stimuli will be delivered to the subjects, and the participants can actively touch a tactile stimulus according to a visual instruction in the MR scanner.

Here, using our MR-compatible system for the delivery of tactile and visual stimuli, we identified a brain region that selectively responds to tactile stimuli through active finger touch compared with visual stimuli. In this system, a tactile stimulus between different types of stimuli can be delivered to the subject by a pneumatic actuator according to a programmed task schedule, and a photosensor detects when a finger touches the tactile stimulus. This pneumatic tactile stimulus delivery system allows the elucidation of brain processes for tactile perception and cognition through natural hand and finger movements.

## Results

### Overview of the pneumatic tactile stimulus delivery system

A tactile object is distinct from a picture or a drawing on the two-dimensional screen as a typical visual object and instead is a volumetric object in the three-dimensional space. Thus, we designed an experimental system to physically deliver this volumetric tactile object specified by a certain texture and shape using an actuator and conveyor. In addition, we considered that unlike most human studies where the participants’ arms and fingers were fixed, a naturalistic condition where the subjects move their fingers freely would be necessary to naturally perceive the tactile stimuli by their fingertips (Hegner et al., 2010; Kitada et al., 2019; Zhang et al., 2005). Thus, the pneumatic tactile stimulus delivery system (pTDS) was designed to deliver a desired stimulus among different tactile stimuli to the subject, allowing active finger touch in the MR scanner (Fig. 1a). In addition, the visual stimuli projected on the screen were used to indicate the time when the subject actively touches the tactile stimulus. With this strategy using both visual and tactile stimuli, in this study, we were able to identify the brain areas that were selectively activated by tactile stimuli compared with visual stimuli.

**Figure 1.**
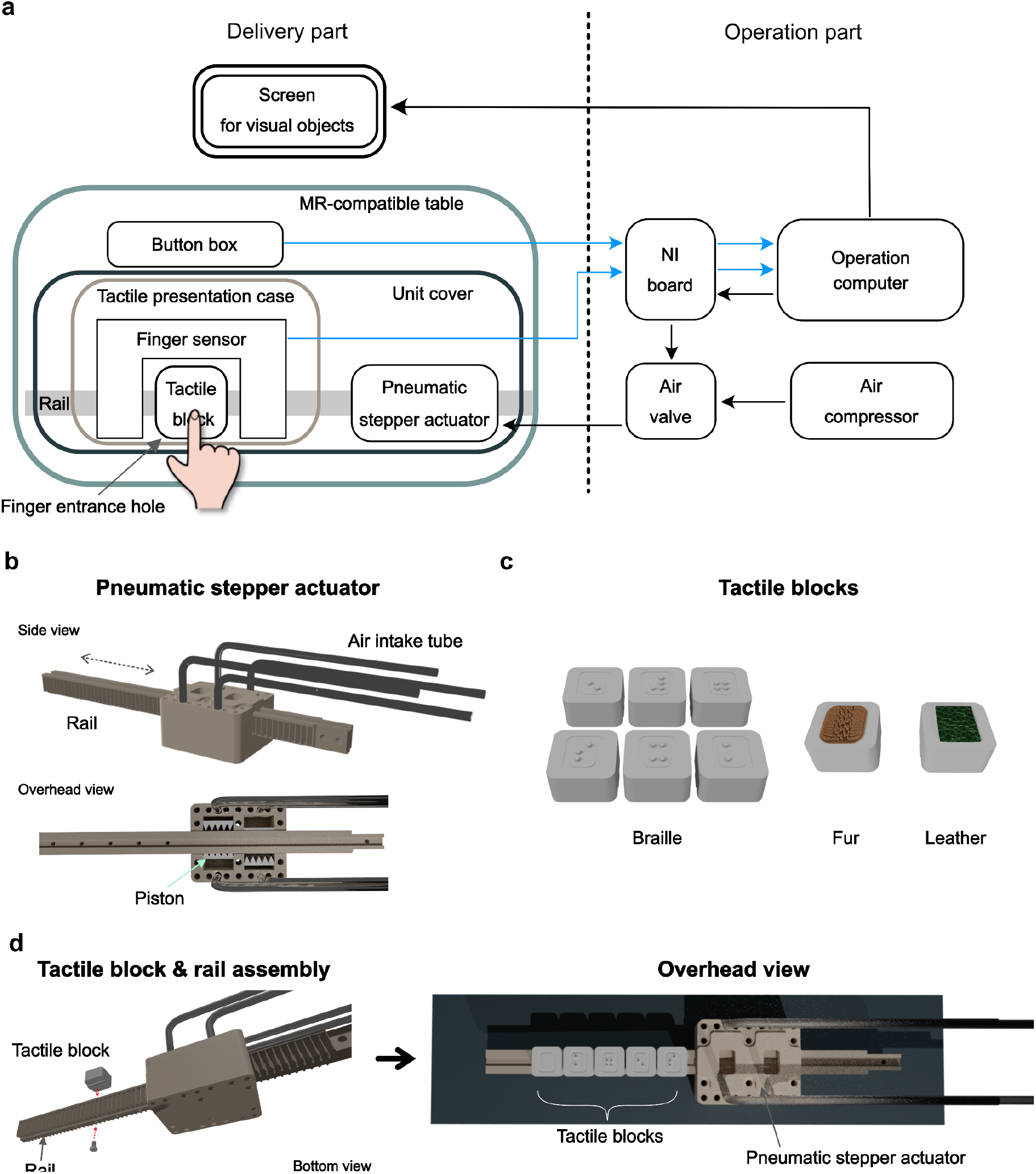
Overview of the pneumatic tactile stimulus delivery system and details of tactile delivery unit with pneumatic actuator and tactile block. (a) Illustration of the overall pneumatic tactile stimulus delivery system (pTDS). The pTDS consists of two components: the operation components that plans and controls the presentation of tactile stimulus, and the delivery component that delivers the stimulus to participants and detects their responses inside the scanner. (b) A 3-D illustration of the pneumatic stepper actuator. The pneumatic stepper actuator is a key component of the pTDS and makes a linear motion with non-electric power. The piston movement by air pressure generates the back-and-forth movement of the rail. (c) An example of tactile blocks. The desired stimulus can be engraved on or attached to the cube-shaped tactile block. (d) Assembly model of the pneumatic stepper actuator with tactile blocks. The pneumatic stepper actuator and the tactile blocks are screwed with plastic bolts (left panel). After assembling the tactile blocks with the rail of the pneumatic stepper actuator, the block can be sent to the desired position through the linear movement of the rail (right panel).

The pTDS largely consists of two components: the operation and delivery component. In the operation component located outside the scanner, a computer plans the presentation of each stimulus in each trial and records behavioral data of finger movements and button presses. In the delivery component located inside the magnet room, the pTDS delivers tactile stimuli and detects when the fingertip touches a tactile stimulus. Details on these two components are explained in the following sections.

### Operation component for pneumatic actuator control and behavioral data acquisition

A computer in the operation component sends information about which tactile block is presented in each trial to the air valves (MHP2-MS1H-5/2-M5, Festo, Germany) that control the pneumatic stepper actuator in the magnet room (Fig. 1a). The computer calculates the number of times the air valves sequentially open to move the pistons and locates a tactile stimulus at the finger entrance hole (Figs. 1b and 2a).

**Figure 2.**
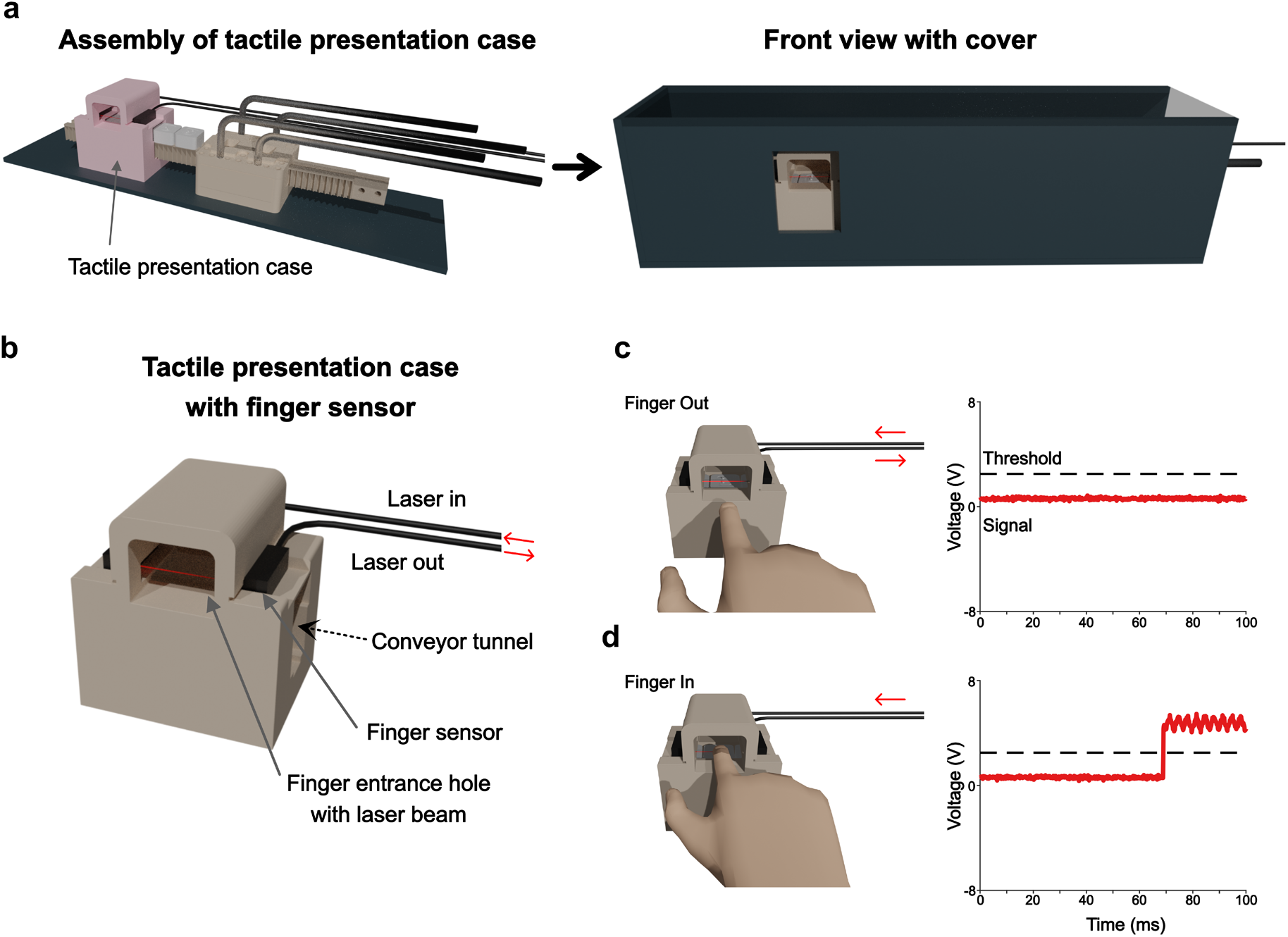
The assembly of the tactile presentation case and mechanism of finger sensor. (a) A 3-D illustration of the assembled tactile presentation case with a pneumatic stepper actuator. The tactile presentation case is fixed on the base plate with plastic bolts (left panel). The unit cover hides the other stimuli except for a tactile block placed in the center of presentation case, allowing the finger to enter (right panel). (b) Detailed description of the tactile presentation case. The finger entrance is detected by a digital laser sensor attached to the side of tactile presentation case. (c–d) Touch detection mechanism with finger sensor. A 0-volt signal is the baseline when the participant’s finger is outside the presentation case (c). When the finger enters the entrance hole and touches the stimulus, the finger sensor sends a 5-volt signal (d).

For the air valve control and behavioral data acquisition, we used a National Instrument (NI) board (PCIe-6353, National Instrument, US) interfaced through a shielded Input/Output connector block (SCB-68, National Instruments) (Fig. 1a). The NI board converts the information of which tactile stimulus is presented into electric signals to control the opening of the air valves. The air valves, which open and close sequentially according to the programmed schedule, control the airflow from the compressor to the pneumatic stepper actuator.

### Software for tactile stimuli presentation and behavioral data acquisition

To deliver the tactile and visual stimuli and record the behavioral data, we used a visual C^++^-based software, Blip (available at www.cocila.net/blip). Blip in the operation computer controls the NI board to send TTL signals for the air valve regulation. In parallel, Blip stores the precise times with event codes of when the stimuli were delivered and when the subjects moved their fingers. For detecting the exact times of touching the tactile stimulus, the oscilloscope function in the Blip software was used to continuously monitor the change of analog input signal (Figs. 2c and d).

### Delivery component for delivering a desired tactile stimulus to the participants

Devices in the delivery component were made from MR-compatible materials to avoid artifacts in the imaging data. There are four main subcomponents: the pneumatic stepper actuator, tactile block, tactile presentation case, and finger sensor (Figs. 1b, 1c, and 2b). Airflow controlled by the air valves moves the teeth of the two pistons in the pneumatic stepper actuator (Fig. 1b) (Groenhuis and Stramigioli, 2018). The sequential up-and-down motions of the piston are converted into the back-and-forth motion of the rail (Fig. 1b). Tactile blocks, shown in Figure 1c, were attached on the rail, and one of the blocks was delivered to the tactile presentation case (Figs. 1d and 2a). Various types of tactile stimuli, such as braille dots, fur, and sandpaper, can be delivered by placing them on the tactile blocks (Fig. 1c). Participants lying in the MR scanner were instructed by a visual stimulus on the screen to touch a tactile stimulus with their fingers through an entrance hole of the presentation case (Figs. 1a and 2d). The finger sensor attached to the presentation case detects the precise times of finger in-and-out movements on the tactile block (Figs. 2b-d).

### Tactile blocks for carrying various types of tactile stimuli

We designed tactile blocks to meet various purposes of tactile research. Different types of tactile stimuli specified by different textures and shapes can be placed on the cubic block (Fig. 1c). The block is made of white polyoxymethylene for MR compatibility and is light enough to be delivered by the pneumatic stepper actuator (4.56 ± 0.02 g). Details on the size and shape of the block are described in Supplementary figure 1.

### Delivering tactile blocks through a rail conveyor of pneumatic stepper actuator

It was challenging to physically deliver the tactile stimuli to the subjects without an electrically powered actuator, which is difficult to use in the MR scanner. We thus used an actuator powered by air pressure as an approach to deliver tactile stimuli (Fig. 1b).

We chose and modified one of the pneumatic stepper actuators (model T-63) that was designed by Groenhuis and Stramigioli because this model has a long rail that can be used as a conveyor for delivering the tactile blocks (Fig. 1b) (Groenhuis and Stramigioli, 2018). This rail can be precisely placed in a specific location and has enough space to hold the tactile blocks, so we adapted this moving rail to convey the tactile stimuli. To mount the tactile blocks on the rail, we tapped screw holes from the top of the rail to the bottom (Fig. 1b, overhead view). Tactile blocks were inserted into a notch of upper part of the rail and secured with plastic screws through the bottom holes (Fig. 1d). The example in Figure 1d shows a rail carrying 5 tactile blocks.

### Tactile presentation case for presenting a desired tactile stimulus and detecting finger movements

We designed a tactile presentation case to deliver a selected tactile stimulus to the participants and examine their finger movements. The tactile presentation case consists of three main components: a conveyor tunnel, finger entrance hole, and finger motion sensor (Figs. 2a and 2b). The tactile blocks on the rail pass through the conveyor tunnel, and a selected tactile stimulus is delivered in the finger entrance hole. This enabled us to present a desired tactile block to the participants.

Laser beam from the optic fiber of a digital laser sensor (FX-505-C2, Panasonic, Japan) placed on both sides of the finger entrance hole detects the participant’s finger movements (Figs. 2b– d). The sensor sends a low voltage baseline signal (approximately 0 volts) when the participant’s finger is out of the entrance hole (Fig. 2c). If the finger is inside the entrance hole and breaks the laser beam on the tactile stimulus, the sensor produces an increased voltage signal (approximately 5 volts) and sends the signal to the operation computer (Fig. 2d). We can set a threshold for the incoming voltage signal using the Blip software (Fig. 2d), while allows us to measure when and how long the participants touch the tactile stimulus. In addition, detecting the signal crossing the threshold can be used to simultaneously control other events, such as turning the visual object presentation on and off.

### MR-compatible tactile delivery system for active finger touch

Brain activity should be examined while the participants are lying on the patient table in MR scanner and minimize their motions. Within the magnet bore, the restriction of movement makes it difficult for participants to find a comfortable height and angle of upper limbs, which may differ depending on the body shape of each individual. Thus, the position of the tactile delivery system should be adjustable for participants to find the most comfortable position for touching the tactile stimuli. We designed a position-adjustable table with a height and angle that are adjustable according to a position where the participant feels comfortable and natural to move their fingers (Fig. 3a). The pneumatic tactile stimulus delivery unit is mounted in the center of the table, and other devices, such as a button box, can be mounted on the rest area of the table (Fig. 3a, overhead view). The table is placed over the participant and fixed to the patient table with plastic bolts and nuts (Fig. 3b, left panel). Using this position-adjustable table, the participants can find comfortable forelimb positions to naturally touch the tactile stimuli and manipulate other devices (Fig. 3b).

**Figure 3.**
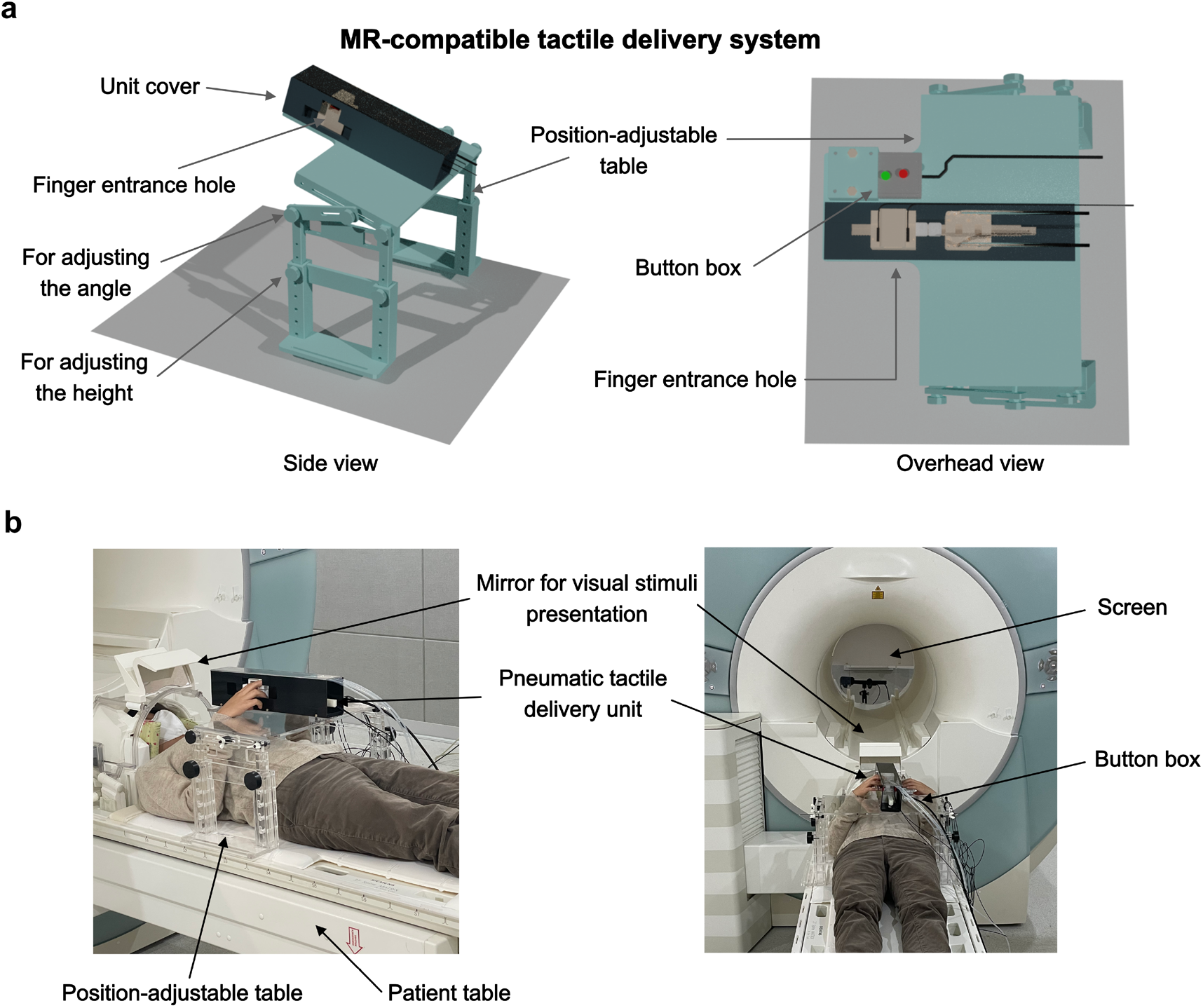
The arrangement of the MR-compatible tactile delivery system for a natural touch of the tactile stimuli with the fingertip. (a) The pneumatic tactile stimulus delivery system with a position-adjustable table. The position-adjustable table with adjustable height and angle enables the individual participants to comfortably and naturally move their fingers in the MR scanner. (b) Photo demonstration of the tactile delivery system with a subject. With the pTDS, participants find their comfortable forelimb positions for natural touch of the tactile stimuli and manipulation of other devices in the MR scanner.

### Tactile stimulus recognition through active finger touch

To test whether participants successfully recognize tactile stimuli in our system compared with visual recognition, we designed a two-modality perception task using braille patterns and fractal images (Fig. 4a). There are two main reasons why we used the braille as tactile stimuli. First, braille is widely used by blind or partially sighted people to read letters with tactile perception, indicating that our tactile perception system can discriminate the patterns of braille dots. Second, 3 x 2 braille cell can produce 63 different combinations of dot patterns, which we can use to systematically test brain processes.

**Figure 4.**
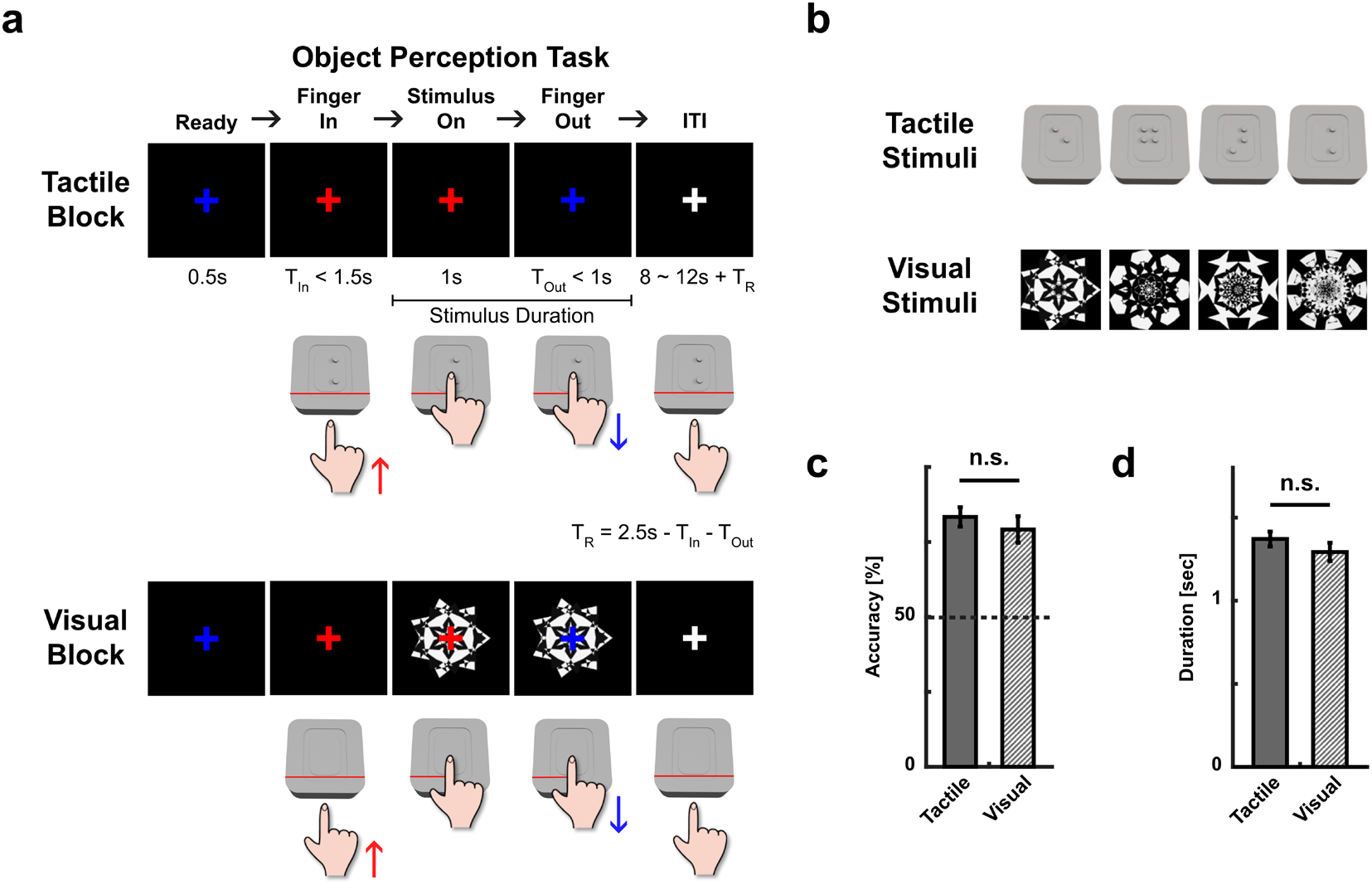
Task paradigm with natural touch and behavioral results. (a) Object perception task consists of two types of blocks: tactile block (upper panel) and visual block (lower panel). During both blocks, participants were asked to insert their right index finger in response to the ‘Finger-In’ cue, which was indicated by the onset of a red fixation cross, and withdraw their index finger in response to the ‘Finger-Out’ cue, which was presented as a blue fixation cross 1 s after the stimulus onset. The presentation of each visual stimulus was turned on when insertion of a participant’s index finger was detected by the light sensor (Stimulus On) and was turned off when the finger was withdrawn. T_in_ is the interval between the onset of the red fixation cross and the time point when light sensor detects the insertion of the finger. The maximum value of T_in_ was 1.5 s. T_out_ is the interval between the onset of the blue fixation cross and the time point when the light sensor detects the absence of the finger in the box. The maximum value of T_out_ was 1 s. If T_in_ or T_out_ is less than its maximum value, the difference between T_in_ (or Tout) and the maximum value, T_R_, was added to ITI. (b) Tactile stimuli (braille patterns) and visual stimuli (fractal images) used in the task. (c) Mean accuracies for the tactile block and visual block. (d) Average duration of each tactile or visual stimulus presentation. n.s., not significant.

In modality perception task, each run of the task consisted of 2 blocks of different modalities: a tactile block and visual block. In the tactile block, the participants were instructed to insert their right index finger in the entrance hole to touch each braille pattern while they viewed a fixation cross at the center of the screen in the absence of any other visual stimulus (Figs. 4a and 4b). In the visual block, they were instructed to view each fractal image presented at the center of the screen while touching a plain surface with their right index finger (Figs. 4a and 4b).

In each trial of both tactile and visual blocks, the participants were instructed to touch a braille stimulus if the blue fixation cross changed to red (‘Finger-In’ cue) and withdraw their finger if the red color of the fixation cross returned to blue (‘Finger-Out’ cue), which was presented 1 s after the stimulus onset (Fig. 4a). To ensure that the participants experience both tactile and visual stimuli for the same amount of time, the visual stimulus was presented while the finger was in the entrance hole.

Both the tactile and visual blocks contained one-back response trials (20% of the total trials), in which the participants had to determine whether the tactile (or visual) stimulus of the current trial matched with the one of the previous trials when they saw a ‘Button-Response’ cue just after ‘Finger-Out’ cue (Fig. S2). We included these test trials to examine whether the participants explicitly recognize the tactile and visual stimuli.

The participants successfully performed the two-modality perception task and showed an accuracy of 83.33 ± 3.23% for the one-back response trials of the tactile blocks and 79.17 ± 4.45% for the one-back response trials of the visual blocks (Fig. 4c). There was no significant difference in the accuracy between the tactile and visual blocks (t_(20)_ = 0.892, p = 0.383), indicating that the participants successfully recognized individual tactile stimuli and that the level of tactile recognition is comparable to that of visual recognition in our system. The mean duration for touching the tactile stimuli was 1.37 ± 0.04 s for tactile blocks and 1.29 ± 0.06 s for visual blocks across the two-modality perception task. There was no significant difference in the duration of stimulus presentation (duration of touch) between the tactile block and visual block (t_(20)_ = 1.053, p = 0.305) (Fig. 4d), indicating comparable duration of stimulus recognition between the tactile and visual blocks.

Our tactile delivery system provides a tool for recognizing individual tactile objects through natural touch during MRI scanning. Furthermore, based on this system, we can compare neural responses elicited during visual and tactile perception under the same conditions except for the sensory modality of experiencing stimuli.

### Sensory modality–dependent cortical responses

Next we investigated neural responses during tactile perception compared with those during visual perception based on our tactile delivery system (pTDS). Specifically, we examined whether early sensory cortical regions are involved during tactile perception compared with visual perception in a modality-specific manner. Prior electrical stimulation studies and neuroimaging studies in humans have shown that tactile perception from fingertip skin engages the middle part of the contralateral somatosensory cortex (Carey et al., 2008; O’Neill et al., 2020; Penfield and Boldrey, 1937; Sanchez Panchuelo et al., 2018), while visual perception involves the early visual cortex in the occipital lobe (Brindley and Lewin, 1968; S A Engel, G H Glover, 1997).

To identify what areas are involved during tactile or visual perception, we first examined the magnitude of BOLD response across the brain during the tactile or visual blocks of the two-modality perception task. We found strong engagement of the left (contralateral) postcentral somatosensory cortex during tactile perception (Fig. 5a, left panel). During visual perception, the involvement of the visual cortex was prominent compared with other regions (Fig. 5a, right panel).

**Figure 5.**
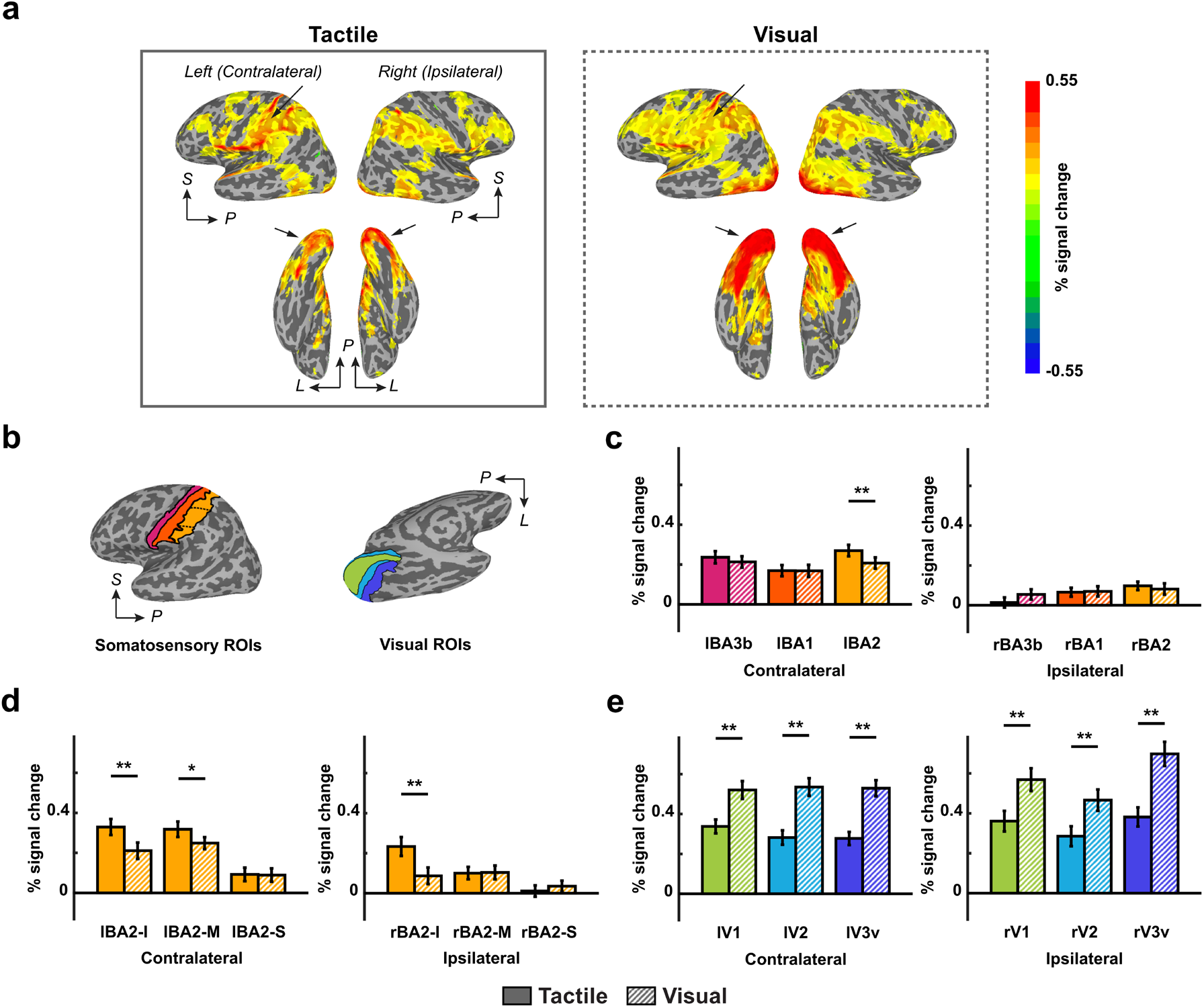
The average magnitude of response during tactile perception and visual perception. (a) Surface maps of brain activation during tactile perception (left) and visual perception (right). The colored areas indicate regions that were significantly activated at stimuli onset of each trial (p < 0.001, uncorrected). S, superior; P, posterior; L, lateral. (b) ROIs mapped on a participant’s inflated left hemisphere. Different ROIs are shown in different colors, and each ROI in (b) and the bar corresponding to the ROI in (c), (d), and (e) are labeled with the same color (magenta: BA3b, orange: BA1, yellow: BA2, light green: V1, light blue: V2, blue: V3v). (c) The averaged magnitude of response across voxels in the left (contralateral) and right (ipsilateral) somatosensory cortical regions (BA3b, BA1 and BA2). (d) The averaged magnitude of response across voxels in left and right BA2 divided into three subregions (I, inferior; M, middle; S, superior). (e) The averaged magnitude of response across voxels in the left and right visual cortical regions (V1, V2, and V3v). *p < 0.05; **p < 0.01; Error bars indicate between-subjects s.e.m.

To further examine the selective involvement cortical regions during tactile or visual perception, we performed ROI analyses focusing on the somatosensory and visual cortex. The neural responses throughout the somatosensory cortex (S1), the Brodmann area 3b (BA3b), BA1, and BA2, were compared between tactile and visual blocks, and a significant difference between the blocks was found in the contralateral BA2 (lBA2) (Fig. 5b). Prior studies have shown that the BA2 region is involved in the integration of tactile and proprioceptive information and is also responsible for shape perception (Bodegård et al., 2001; Keysers et al., 2010). Indeed, we observed greater response in the left BA2 (lBA2) during the tactile perception than during the visual perception (t_(20)_ = 2.868, p = 0.010) (Fig. 5c). Given that the middle part of the somatosensory cortex is mainly engaged in sensation from hand and fingers (O’Neill et al., 2020; Sanchez Panchuelo et al., 2018), we divided BA2 of each hemisphere into three parts along the superior-inferior axis. Consistent with the results observed in the whole brain responses (Fig. 5a), we found a significantly greater response magnitude during tactile perception than during visual perception in the middle part of the left (contralateral) BA2 (lBA2-M) (t_(20)_ = 2.286, p = 0.033) (Fig. 5d). This difference between the tactile and visual perception blocks was not observed in the right (ipsilateral) BA2-M (rBA2-M) (t_(20)_ = − 0.104, p = 0.919). Moreover, stronger responses were observed in lBA2-M than rBA2-M during tactile perception (t_(20)_ = 6.894, p < 0.001). Greater responses during tactile perception than visual perception were also found in the inferior part of the bilateral BA2 (lBA2-I, rBA2-I) (left: t_(20)_ = 5.153, p < 0.001; right: t_(20)_ = 2.967, p = 0.008), and no significant difference between lBA2-I and rBA2-I was found (t_(20)_ = 1.730, p = 0.099). Unlike the results for the middle and inferior part of BA2, there was no significant difference between the modalities in the superior part of BA2 (lBA2-S: t_(20)_ = 0.117, p = 0.908; rBA2-S: t_(20)_ = −0.809, p = 0.428). These results show tactile-modality specific responses in the somatosensory cortex despite the different responses between subregions of the somatosensory cortex.

While greater responses were observed during tactile perception than visual perception in the somatosensory cortex, the opposite tendency was observed in early visual regions. We found significantly stronger responses during the visual blocks compared with the tactile blocks in bilateral V1, V2, and V3 (lV1: t_(20)_ = −6.752, p < 0.001; rV1: t_(20)_ = −7.000, p < 0.001; lV2: t_(20)_ = −11.781, p < 0.001; rV2: t_(20)_ = −7.340, p < 0.001; lV3v: t_(20)_ = −9.840, p < 0.001; rV3: t_(20)_ = −9.455, p < 0.001) (Fig. 5e).

We confirmed that lBA2-M and early visual regions (V1, V2, and V3v) showed a significantly different tendency of responses during tactile or visual perception based on two-way repeated measures ANOVA with ROI and sensory modality (tactile and visual) as within-subject factors. We found a significant interaction effect for lBA2-M and each of the visual ROIs (lV1: F_(1,20)_ = 44.500, p < 0.001; lV2: F_(1,20)_ = 106.502, p < 0.001; lV3v: F_(1,20)_ = 118.907, p < 0.001; rV1: F_(1,20)_ = 43.476, p < 0.001; rV2: F_(1,20)_ = 42.216, p < 0.001; rV3v: F_(1,20)_ = 93.248, p < 0.001). Taken together, these results confirm that our MR-compatible tactile stimulation system elicits tactile perception-specific responses in the human sensory cortex.

## Discussion

We developed a tactile stimulus delivery system, pTDS, that provides tactile stimuli on a programmed schedule in an MR-compatible manner, in which participants actively touch stimuli while their brain activities are monitored in the MRI scanner. This MR-compatible system based on a pneumatic actuator and a photosensor allows the comparison of different conditions of active touch, enabling systematic investigation of neural processes during the perception of individual tactile stimuli. Using the pTDS, we successfully demonstrated the tactile modality-selective brain responses elicited by voluntary finger touch, indicating the feasibility and usefulness of pTDS in human fMRI studies.

### Tactile pattern recognition by active touching

We demonstrated the utility of the newly designed tactile stimuli delivery system by measuring modality-specific neural responses during perception of tactile and visual stimuli. Using the pTDS, we could implement the object perception task in which participants recognized tactile or visual stimuli and responded to intermittent 1-back task. Participants touched the tactile objects (braille patterns or blank surface) by voluntarily moving their finger into the presentation case and exploring the object surface to recognize the shape. In prior studies of tactile perception, participants’ voluntary movements were restricted and they were allowed to perceive only given stimuli with hands and figures in a fixed position (Hegner et al., 2010; Kitada et al., 2019; Zhang et al., 2005). However, the pTDS allows us to capture the way humans interact with objects by their hands in a more natural manner, as humans often figure out identity, shape, or size of objects through tactile sensation by voluntarily reaching and actively exploring the surface of the structure. Investigation of the neural processing that underlies active tactile perception is essential for understanding the tactile processes in the brain, as the neural circuitry of active and passive tactile perception may differ in that active tactile perception involves integration of sensory and motor information. Indeed, previous studies support the dissociation of the two processes in the brain (Kavounoudias et al., 2008; Simões-Franklin et al., 2011; Valenza et al., 2001). While modeling the naturalistic behavior of tactile shape recognition, timing and duration of touch need to be controlled for practical reasons to conduct neuroimaging experiments. In classical methods, stimuli can be presented and removed by an experimenter in the MR scanner in response to auditory cues through headphones (Grefkes et al., 2002). We presented a visual cue (color change of the fixation cross) to participants to guide the timing of voluntary insert and withdrawal of their finger. Participants successfully recognized the braille patterns as much as the visual fractal objects. The accuracy for the 1-back task and the tactile stimulus duration, which is determined by reaction time to the withdrawal cue (blue fixation cross), were both comparable between tactile and visual condition. These results indicate that the given time (1 s) between the finger-in and presentation of the finger withdrawal cue was sufficient for both the tactile perception and the visual perception.

### Modality-specific neural responses for tactile pattern recognition

Consistent with the prior studies reporting neural substrates of tactile signal processes in the brain (Carey et al., 2008; O’Neill et al., 2020; Penfield and Boldrey, 1937; Sanchez Panchuelo et al., 2018), we found modality-specific neural responses in the contralateral somatosensory cortex during the tactile braille pattern recognition. In particular, significantly stronger activation was found especially in the contralateral BA2 during the tactile perception compared with visual perception (Fig. 5c). We carefully designed the two-modality perception task to examine the neural responses induced by tactile recognition of braille stimuli rather than the responses related to motor control or sensation of mere contact of the finger. The participants were instructed to touch the blank surface while they viewed the visual images, even during visual blocks of the task. The comparable level of activation in BA3b and BA1 between the tactile and visual blocks (Fig. 5c) may reflect the effect of mere contact of the finger, which was commonly involved in both blocks. Given that BA2 has been considered to be involved in the processing of integrated tactile information and shape recognition (Bodegård et al., 2001; Keysers et al., 2010), the stronger responses observed in BA2 during tactile blocks than during visual blocks of the two-modality perception task may reflect neural processing of braille pattern–specific tactile recognition.

We further compared the neural responses in detail by dividing the BA2 along the longitudinal axis (Fig. 5d). According to the prior electrical stimulation (Penfield and Boldrey, 1937) and neuroimaging studies (O’Neill et al., 2020; Sanchez Panchuelo et al., 2018) focusing on the human somatosensory cortex, the tactile sensation from the digits is mainly associated with the middle part of the primary somatosensory cortex while the sensation from the face and foot is mapped on the inferior part and the superior (medial) part, respectively. Consistent with these results, we found that the neural responses during braille perception with the index finger were prominent in the middle part of the contralateral BA2. We additionally observed greater responses during tactile blocks compared with visual blocks in the inferior BA2 of the bilateral hemispheres. The tactile perception–specific responses in this region may possibly reflect higher level tactile processing given the close location of this region to the secondary somatosensory cortex in the parietal operculum (Penfield and Jasper, 1954; Woolsey et al., 1979). Indeed, the bilateral secondary somatosensory cortex has been reported to be activated when tactile stimulation was given to right hand (Lamp et al., 2019). These observations of differential neural responses in the known neural substrates for the tactile perception suggest that the system we developed can be used in fMRI experiments to investigate the neural mechanism of cognitive processes in the tactile domain.

### Other potential scalabilities and implications of the pTDS

The pTDS has many implications for future fMRI studies using tactile modality. First, the pTDS is designed to deliver various stimuli as needed. Since there have been attempts to study numerous tactile stimuli using fMRI, the pTDS can be an advantage to examine the various properties of tactile stimuli (Hegner et al., 2010; Kitada et al., 2019; Lee et al., 2019; Podrebarac et al., 2014; Zhang et al., 2005). The number of tactile stimuli delivered by the pTDS can be increased by using a longer rail holding more tactile blocks. This approach allows testing and comparison of different properties of tactile stimuli simultaneously.

The pTDS is also applicable in more complex experimental settings, such as a choice task using multiple tactile stimuli. For example, two sets of the pTDS may be used for delivering two different tactile stimuli at the same time. The two pTDSs can be positioned in parallel on the adjustable table, allowing the participant to sequentially touch and select one of the two.

Another advantage of the pTDS is its compatibility with other fMRI devices such as an eye-tracking system, screen, and button box. Depending on the study goal, the researchers can place other devices such as a joystick and a button box on the position-adjustable table together with the pTDS. This allows the study of neural representation of various modalities simultaneously with tactile modality.

## Methods

### Participants

Twenty-one neurologically intact right-handed participants (mean age 24.48 ± 3.63 years, range 20–33, 10 females) took part in the experiment. The participants reported that they had normal or corrected-to-normal vision. All participants provided informed consent for the procedure. The experimental procedure was approved by the Institutional Review Board of Seoul National University, and Korea Advanced Institute of Science and Technology.

### Visual stimuli

We used 4 visual fractal objects created using Fractal Geometry (Kang et al., 2021; Miyashita et al., 1991) (Fig. 4b). The mean luminance was equalized across the images using Spectrum, Histogram, and Intensity Normalization and Equalization (SHINE) toolbox written with MATLAB (www.mapageweb.umontreal.ca/gosselif/shine). Grayscale images of the fractal objects were used to minimize the color effect on brain response.

### Tactile stimuli

We used 4 different braille patterns engraved on top of each polyoxymethylene block (Fig. 4b). The size of each tactile block was 18 × 18 × 12 mm and the diameter and height of each dot was 2 × 1 mm (an example tactile block in shown in Fig. S1).

### Object Perception Task

We designed a two-modality perception task using braille patterns and fractal images (Fig. 4a). Each run of the task consisted of two blocks of different modalities: a tactile block and visual block. In the tactile block, the participants were instructed to insert their right index finger in the entrance hole to touch each braille pattern while they viewed a fixation cross at the center of the screen in absence of any other visual stimulus (Figs. 4a and 4b). In the visual block, the participants were instructed to view each fractal image presented at the center of the screen while touching the plain surface with their right index finger (Figs. 4a and 4b). The finger motions of participants were the same in both blocks to minimize the motion-induced MR responses.

In each trial with the tactile and visual blocks, the participants were asked to touch a braille stimulus in the hole when the blue fixation cross changed to red (‘Finger-In’ cue) and withdraw their finger from the hole when the red color of the fixation cross returned blue (‘Finger-Out’ cue), which was presented 1 s after the stimulus onset (Fig. 4a). To ensure that the participants experienced both tactile and visual stimuli for the same amount of time, the visual stimulus was presented only while the finger was in the entrance hole.

Both the tactile and visual blocks contained one-back response trials (20 % of the total trials), in which the participants had to respond whether the tactile (or visual) stimulus of the current trial was the same as the one from the previous trial when they saw a ‘Button-Response’ cue just after ‘Finger-Out’ cue (Fig. S2). These test trials were used to examine whether the participants explicitly recognize the tactile and visual stimuli.

### fMRI Acquisition

Participants were scanned on a 3T Siemens MAGNETOM Trio located in the Seoul National University. Echo-planar imaging (EPI) data were acquired using a 32-channel head coil, with an in-plane resolution of 2.8 mm x 2.8 mm, and 40 2.5 mm slices (0.25 mm inter-slice gap, repetition time (TR) = 2000 ms, echo time (TE) = 25 ms, matrix size 76 x 76, field of view (FOV) = 210 mm). Whole brain volumes were scanned, and slices were oriented approximately parallel to the base of the temporal lobe. Anatomical images were scanned with standard magnetization-prepared rapid-acquisition gradient echo (MPRAGE) sequence after the experimental runs of the fMRI session.

### Regions-of-interest (ROIs)

For ROI-based analyses, somatosensory and early visual regions were automatically defined by parcellation of FreeSurfer. Brodmann area 3b, 1, and 2 (BA3b, BA1, and BA2, respectively) were first derived based on ‘BA3b_exvivo.label,’ ‘BA1_exvivo.label,’ and ‘BA2_exvivo.label,’ respectively and subsequently modified by excluding overlapped areas between them; overlap with BA3a (derived from ‘BA3a_exvivo.label’) was excluded from BA3b, overlap with BA3b was excluded from BA1, and overlap with BA1 was excluded from BA2. For the detailed analyses, BA2 of each hemisphere was divided into three parts with equal length along the longitudinal axis (I: inferior, M: middle, S: superior). We used vcAtlas labels to define areas in the ventral visual stream (Rosenke et al., 2018): ‘hOc1.mpm.vpnl.label’ for V1, ‘hOc2.mpm.vpnl.label’ for V2, and ‘hOc3v.mpm.vpnl.label’ for V3v.

### fMRI Data Analysis

fMRI data analysis was conducted using AFNI (http://afni.nimh.nih.gov/), SUMA (AFNI surface mapper), FreeSurfer, and custom MATLAB scripts. Data preprocessing including slice-time correction and motion correction was conducted.

To derive the event-related response magnitudes of BOLD signal during tactile or visual perception, we generated a standard general linear model using AFNI software package (GAM function of 3dDeconvolve). The β-value of each voxel was derived for the onset of each stimulus presentation and was normalized by the average magnitude of the response for each run (percent signal change). For whole brain map of the activation level, average value of percent signal change across the voxels within each searchlight sphere (radius 9.8 mm, corresponding to ~123 voxels) was assigned to each centered voxel to smooth the data to derive group averaged surface map.

To test temporal signal-to-noise-ratio (tSNR) of the EPI data acquired in the presence of our tactile delivery system, we calculated tSNR for each voxel as the mean value of the time series divided by standard deviation of the signal. The average tSNR across all voxels was 54.021 (averaged across subjects, SEM 2.332), which is within the normal range (Murphy et al., 2007).

### Statistical Analysis

We compared behavioral data and brain activation level between tactile and visual modality mainly using paired t-test (two-tailed). Activation levels were compared to basal level (zero) by one-sample t-test (two-tailed). We also used two-way repeated measures ANOVA (tests of within-subjects effects) to examine the effect of interaction between the modalities and the ROIs. We used SPSS and MATLAB for statistical analyses.

## Data and Code Availability

The experimental code and software used in this study are available at this link (www.cocila.net/blip). The behavioral and neuroimaging data are not able to be made openly available because of the need for approval from the Institutional Review Boards. Sharing and reuse of data require a reasonable request by sending an e-mail to the corresponding author with the name of P.I., affiliation, and purpose of the study, and approval from the Institutional Review Boards.

## ACKNOWLEDGMENTS

This work was supported by the Neurological Disorder Research Program (NRF-2020M3E5D9079908) and the Basic Science Research Program (NRF-2019R1A2C2005213 and NRF-2020R1A2C2007770) through the National Research Foundation (NRF) of Korea, and Creative-Pioneering Researchers Program through Seoul National University.

## Supplementary figure legends

**Supplementary Figure 1.**
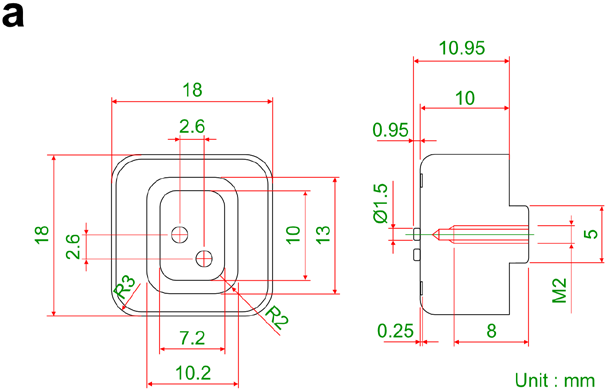
Details of the tactile block. Drawing of a single tactile block with an engraved braille pattern. An 8-mm-long screw hole was made under the cube block for insertion into a notch in the upper part of the rail.

**Supplementary Figure 2.**
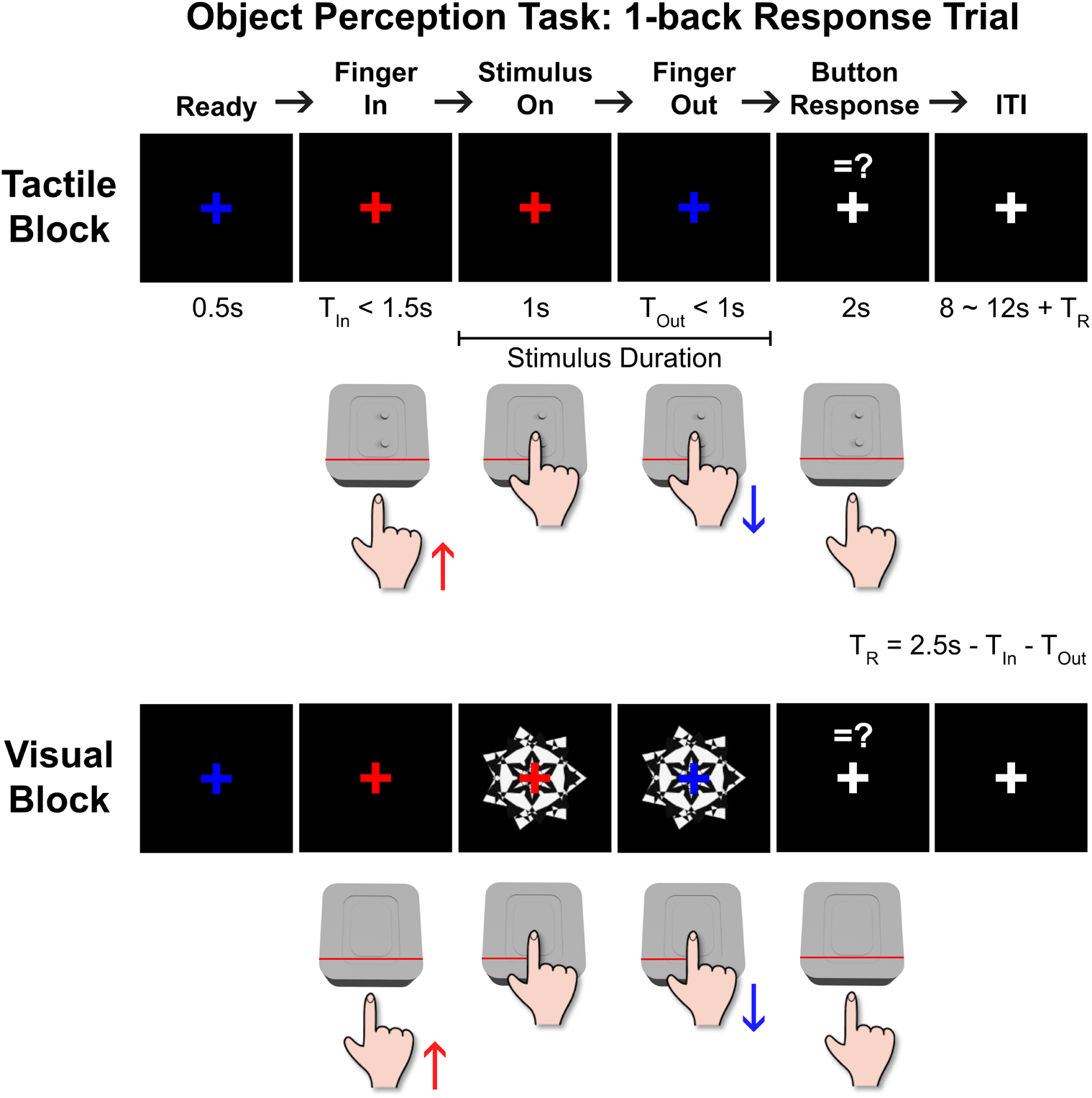
Task paradigm for 1-back response trials. Both the tactile and visual blocks contained 1-back response trials (20 % of the total trials), in which participants had to respond whether the object of the current trial was the same with the one of the previous (1-back) trial. In these trials, the ‘Button-Response’ cue was presented just after the finger was withdrawn, and the participants pressed the button if the matched object was presented.

## References

Bodegård, A., Geyer, S., Grefkes, C., Zilles, K., Roland, P.E., 2001. Hierarchical processing of tactile shape in the human brain. Neuron 31, 317–328. 10.1016/S0896-6273(01)00362-2

Brindley, B.Y.G.S., Lewin, W.S., 1968. The sensations produced by electrical stimulation of the visual cortex. J. Physiol. 196, 479–493.

Carey, L.M., Abbott, D.F., Egan, G.F., Donnan, G.A., 2008. Reproducible activation in BA2, 1 and 3b associated with texture discrimination in healthy volunteers over time. Neuroimage 39, 40–51. 10.1016/j.neuroimage.2007.08.026

Debowska, W., Wolak, T., Soluch, P., Orzechowski, M., Kossut, M., 2013. Design and evaluation of an innovative MRI-compatible Braille stimulator with high spatial and temporal resolution. J. Neurosci. Methods 213, 32–38. 10.1016/j.jneumeth.2012.12.002

Grefkes, C., Weiss, P.H., Zilles, K., Fink, G.R., 2002. Crossmodal processing of object features in human anterior intraparietal cortex: an fMRI study implies equivalencies between humans and monkeys. Neuron 35, 173–184.

Groenhuis, V., Stramigioli, S., 2018. Rapid Prototyping High-Performance MR Safe Pneumatic Stepper Motors. IEEE/ASME Trans. Mechatronics 23, 1843–1853. 10.1109/TMECH.2018.2840682

Hegner, Y.L., Lee, Y., Grodd, W., Braun, C., 2010. Comparing tactile pattern and vibrotactile frequency discrimination: A human fMRI study. J. Neurophysiol. 103, 3115–3122. 10.1152/jn.00940.2009

Kang, J., Kim, H., Hwang, S.H., Han, M., Lee, S.H., Kim, H.F., 2021. Primate ventral striatum maintains neural representations of the value of previously rewarded objects for habitual seeking. Nat. Commun. 12, 1–13. 10.1038/s41467-021-22335-5

Kavounoudias, A., Roll, J.P., Anton, J.L., Nazarian, B., Roth, M., Roll, R., 2008. Proprio-tactile integration for kinesthetic perception: an fMRI study. Neuropsychologia 46, 567–575.

Keysers, C., Kaas, J.H., Gazzola, V., 2010. Somatosensation in social perception. Nat. Rev. Neurosci. 11, 417–428. 10.1038/nrn2833

Kitada, R., Doizaki, R., Kwon, J., Tanigawa, T., Nakagawa, E., Kochiyama, T., Kajimoto, H., Sakamoto, M., Sadato, N., 2019. Brain networks underlying tactile softness perception: A functional magnetic resonance imaging study. Neuroimage 197, 156–166. 10.1016/j.neuroimage.2019.04.044

Lamp, G., Goodin, P., Palmer, S., Low, E., Barutchu, A., Carey, L.M., 2019. Activation of bilateral secondary somatosensory cortex with right hand touch stimulation: A meta-analysis of functional neuroimaging studies. Front. Neurol. 10, 1–14. 10.3389/fneur.2018.01129

Lee, H., Lee, E., Jung, J., Kim, J., 2019. Surface Stickiness Perception by Auditory, Tactile, and Visual Cues. Front. Psychol. 10, 1–8. 10.3389/fpsyg.2019.02135

Middlebrooks, E.H., Frost, C.J., Tuna, I.S., Schmalfuss, I.M., Rahman, M., Old Crow, A., 2017. Reduction of motion artifacts and noise using independent component analysis in task-based functional MRI for preoperative planning in patients with brain tumor. Am. J. Neuroradiol. 38, 336–342. 10.3174/ajnr.A4996

Miyashita, Y., Higuchi, S.I., Sakai, K., Masui, N., 1991. Generation of fractal patterns for probing the visual memory. Neurosci. Res. 12, 307–311. 10.1016/0168-0102(91)90121-E

Murphy, K., Bodurka, J., Bandettini, P.A., 2007. How long to scan? The relationship between fMRI temporal signal to noise ratio and necessary scan duration. Neuroimage 34, 565–574. 10.1016/j.neuroimage.2006.09.032

O’Neill, G.C., Sengupta, A., Asghar, M., Barratt, E.L., Besle, J., Schluppeck, D., Francis, S.T., Sanchez Panchuelo, R.M., 2020. A probabilistic atlas of finger dominance in the primary somatosensory cortex. Neuroimage 217, 116880. 10.1016/j.neuroimage.2020.116880

Penfield, W., Boldrey, E., 1937. Somatic motor and sensory representation in the cerebral cortex of man as studied by electrical stimulation. Brain 60, 389–443.

Penfield, W., Jasper, H., 1954. Epilepsy and the functional anatomy of the human brain.

Podrebarac, S.K., Goodale, M.A., Snow, J.C., 2014. Are visual texture-selective areas recruited during haptic texture discrimination? Neuroimage 94, 129–137. 10.1016/j.neuroimage.2014.03.013

Rosenke, M., Weiner, K.S., Barnett, M.A., Zilles, K., Amunts, K., Goebel, R., Grill-Spector, K., 2018. A cross-validated cytoarchitectonic atlas of the human ventral visual stream. Neuroimage 170, 257–270. 10.1016/j.neuroimage.2017.02.040

S A Engel, G H Glover, B.A.W., 1997. Retinotopic organization in human visual cortex and the spatial precision of functional MRI. Cereb. cortex 7, 181–192.

Sanchez Panchuelo, R.M., Besle, J., Schluppeck, D., Humberstone, M., Francis, S., 2018. Somatotopy in the Human Somatosensory System. Front. Hum. Neurosci. 12, 1–14. 10.3389/fnhum.2018.00235

Simões-Franklin, C., Whitaker, T.A., Newell, F.N., 2011. Active and passive touch differentially activate somatosensory cortex in texture perception. Hum. Brain Mapp. 32, 1067–1080.

Valenza, N., Ptak, R., Zimine, I., Badan, M., Lazeyras, F., Schnider, A., 2001. Dissociated active and passive tactile shape recognition: a case study of pure tactile apraxia. Brain 124, 2287–2298.

Woolsey, C.N., Erickson, T.C., Gilson, W.E., 1979. Localization in somatic sensory and motor areas of human cerebral cortex as determined by direct recording of evoked potentials and electrical stimulation. J. Neurosurg. 51, 476–506.

Zhang, M., Mariola, E., Stilla, R., Stoesz, M., Mao, H., Hu, X., Sathian, K., 2005. Tactile discrimination of grating orientation: fMRI activation patterns. Hum. Brain Mapp. 25, 370–377. 10.1002/hbm.20107

